# A Broadly Conserved Protective Epitope on the Lyme Disease Vaccine Antigen, OspA

**DOI:** 10.64898/2025.12.24.696337

**Authors:** Graham G. Willsey, Michael J. Rudolph, Carol Lyn Piazza, Yang Chen, Grace Freeman-Gallant, Lisa A. Cavacini, David J. Vance, Nicholas J. Mantis

**Affiliations:** Division of Infectious Diseases, Wadsworth Center, New York State Department of Health, Albany, NY 12208; New York Structural Biology Center, New York, NY 10027; Department of Medicine, University of Massachusetts Chan School of Medicine, Worcester, MA 01655

## Abstract

Lyme disease, caused by the spirochete, *Borreliella burgdorferi* sensu latu (Bbsl), is a debilitating tickborne infection of increasing incidence in North America, Europe and Asia. While vaccines based on Outer surface protein A (OspA) have proven highly efficacious at blocking Bbsl tick-to-human transmission, the high degree of antigenic variability among the major OspA serotypes (ST) has made the development of a broadly cross protective vaccine difficult. Recent profiling of protective human monoclonal antibodies (mAbs) has suggested the existence of conserved epitopes situated within OspA’s central β-sheet (CBS), although direct comparisons of cross-serotype functionality has been hindered by biological differences among the major Bbsl genospecies. To address these issues, we developed a panel of isogenic *B. burgdorferi* viability reporter strains expressing the seven major OspA serotypes (ST1-7) and probed them with CBS-targeting mAbs to evaluate their complement-dependent borreliacidal activity. The mAbs segregated into three distinct classes with varying degrees of borreliacidal activity: class 1 mAbs exhibited potent killing against all seven OspA serotypes, while classes 2 and 3 had restricted or no activity against two of the seven serotypes. Structural analysis of Fabs derived from each class of mAbs in complex with OspA ST1 showed that they target overlapping epitopes spanning β-strands 6-10 and involve contact with largely invariant residues. Further analysis of *B. burgdorferi* reporter strains expressing OspA variants from 17 additional Bbsl genospecies identified Lys-107 as a determinant of susceptibility for nearly all CBS mAbs. Taken together, these findings raise the prospect of structure-based design of a broadly protective monovalent Lyme disease vaccine.

**AUTHOR SUMMARY:** Lyme disease, caused by the spirochete, *Borreliella burgdorferi* sensu latu (Bbsl), is a potentially debilitating tickborne infection of increasing incidence in North America, Europe and Asia. While vaccines based on Outer surface protein A (OspA) have proven highly efficacious at blocking Bbsl tick-to-human transmission, the genetic and serological heterogeneity of OspA across Borrelia genospecies has complicated matters. In this report, we use a collection of transmission-blocking human monoclonal antibodies to delineate a protective region (epitope) within the central core of OspA that is conserved across all major OspA serotypes. These results have important implications for engineering a broadly reactive Lyme disease vaccine.

## INTRODUCTION

The spirochete *Borreliella burgdorferi* sensu stricto (Bbss) is the primary etiologic agent of Lyme disease, the most common tick-borne infection in the United States (1). The disease typically presents with a characteristic “bullseye” rash (erythema migrans) before potentially progressing to a disseminated infection, which may involve carditis, neuroborreliosis, and/or Lyme arthritis (2). Although antibiotic treatment is generally effective at any stage of infection, a subset of individuals experiences persistent symptoms known as post-treatment Lyme disease (PTLD) syndrome (3). In Europe and Asia, Lyme disease is also caused by three additional genospecies within the *Borreliella burgdorferi* sensu lato (Bbsl) complex: *B. afzelii*, *B. garinii*, and *B. bavariensis*. While these four genospecies share key pathogenic traits, they are genetically and antigenically heterogeneous and exhibit distinct clinical manifestations. This heterogeneity has posed significant challenges to the development of broadly protective Lyme disease vaccines suitable for use across North America, Europe, and Asia.

The only Lyme disease vaccine licensed for human use in the United States, LYMERix, was based on outer surface protein A (OspA) serotype 1 (ST1), the predominant OspA ST expressed by *B. burgdorferi* in North America (4, 5). OspA is a ∼31 kDa lipoprotein abundantly expressed on the surface of *B. burgdorferi* in the midgut of unfed ticks. During feeding, OspA expression is downregulated as the spirochete migrates through the hemolymph and salivary glands. Nonetheless, transmission is effectively blocked if a tick feeds on a host with pre-existing OspA antibodies induced by vaccination (6–10).

Similarly, administration of certain mouse or human monoclonal antibodies (mAbs) targeting OspA prior to tick challenge is sufficient to protect rodents and non-human primates from *B. burgdorferi* infection (7, 11, 12). While the precise mechanisms by which these antibodies block transmission are not fully understood, evidence supports a model in which they engage spirochetes within the tick midgut (10, 13).

As the incidence of Lyme disease continues to rise in both the United States and Europe, there is renewed interest in developing OspA-based vaccines. To provide broad protection beyond North America, such vaccines must target not only *B. burgdorferi* (ST1), but also six additional OspA serotypes (ST2–7) associated with *B. afzelii* (ST2), *B. garinii* (ST3, ST5–7), and *B. bavariensis* (ST4) (14, 15). Research dating back over three decades has shown that OspA-based immunization typically confers strong protection against homologous serotype challenge, but limited or no protection against heterologous serotypes (16–18).

These findings, combined with the identification of immunodominant, highly protective B cell epitopes localized to OspA’s C-terminal region, have driven the design and evaluation of multivalent OspA vaccines (9, 19–22). One such candidate, VLA15, incorporates chimeric heterodimers of the C-terminal third of OspA ST1–6 and is currently in Phase III clinical trials (23–26). However, reports of cross-serotype immunity following immunization with recombinant OspA suggest the presence of at least some conserved protective epitopes (17), which could have important implications for future vaccine design.

Monoclonal antibodies (mAbs) have been instrumental in identifying cross-protective epitopes on highly variable surface antigens from pathogens such as influenza virus hemagglutinin (27, 28), HIV-1 envelope glycoprotein (29), and *Plasmodium* species (30), among others (31). In the context of Lyme disease, several human mAbs have been shown to mediate complement-dependent borreliacidal activity against OspA from *B. burgdorferi* (ST1), *B. afzelii* (ST2), and *B. bavariensis* (ST4) (11). Notably, two such mAbs - 221-7 and 857-2 - have demonstrated the ability to block tick-mediated transmission of *B. burgdorferi* (ST1) in a murine model, with 221-7 also shown to be protective in a non-human primate model (11, 12). Epitope mapping via hydrogen-deuterium exchange mass spectrometry (HDX-MS) localized the targets of these cross-reactive mAbs to overlapping regions within OspA’s central β-sheet (CBS), a relatively conserved domain compared to the more variable C-terminus (11, 14, 32). Structural analysis of mAb 221-7—currently under consideration as a pre-exposure prophylactic—revealed an epitope potentially associated with broad reactivity, although its full breadth of binding and borreliacidal activity had not yet been comprehensively characterized (12). In this study, we address this gap by using a panel of human OspA-specific mAbs to identify conserved, protective epitopes across all major OspA serotypes associated with Lyme disease causing spirochetes.

## RESULTS

Differences in *Bbsl* genospecies growth rates, intrinsic serum sensitivities, and susceptibility to commercially available complement sources have hindered the development of a standardized OspA binding and complement-dependent bactericidal assay suitable for all Bbsl serotypes and in silico types (ISTs) (14, 15, 23, 25, 33). To resolve this issue, we constructed a panel of isogenic *B. burgdorferi* reporter strains expressing each of the seven major OspA serotypes (OspA_ST1–7_). This was accomplished by modifying an existing IPTG-inducible *mScarlet-I* viability reporter plasmid (pGW189) to encode different *ospA* alleles under the control of the native *ospAB* promoter from *B. burgdorferi* B31 (34). Reporter plasmids encoding *ospA* alleles from *B. burgdorferi* B31 (ST1), *B. afzelii* PKo (ST2), *B. garinii* PBr (ST3), *B. bavariensis* PBi (ST4), *B. garinii* PHei (ST5), *B. garinii* DK29 (ST6), and *B. garinii* T25 (ST7) were introduced into *B. burgdorferi* strain HB19-R1, a high passage North American isolate that lacks *ospA* due to the loss of lp54 (**Table S1-S2**) (35, 36).

To validate the OspA reporter strains, we assessed the reactivity of two mAbs, 857-2 and LA-2, against the panel and compared their profiles to corresponding native Bbsl strains (**Figure 1**; **Figure S1**). 857-2 was reactive with all seven OspA serotypes expressed in the HB19-R1 background, with median fluorescence intensity (MFI) values ranging from 2500-6400. In contrast, LA-2 was only reactive with OspA_ST1_ (>70% reactivity; MFI>7,000). Neither 857-2 nor LA-2 reacted with HB19-R1 carrying the empty vector (MFI<55). The same OspA reactivity profiles were observed when 857-2 and LA-2 were used to probe the native Bbsl strains (**Figure S1**): 857-2 recognized all seven OspA serotypes, while LA-2 only recognized OspA_ST1_. These results not only demonstrate that the HB19-R1 reporter strains are reliable proxies for assessing antibody binding to different OspA serotypes but also confirm that 857-2 can recognize the seven major OspA serotypes (34).

**Fig. 1.**
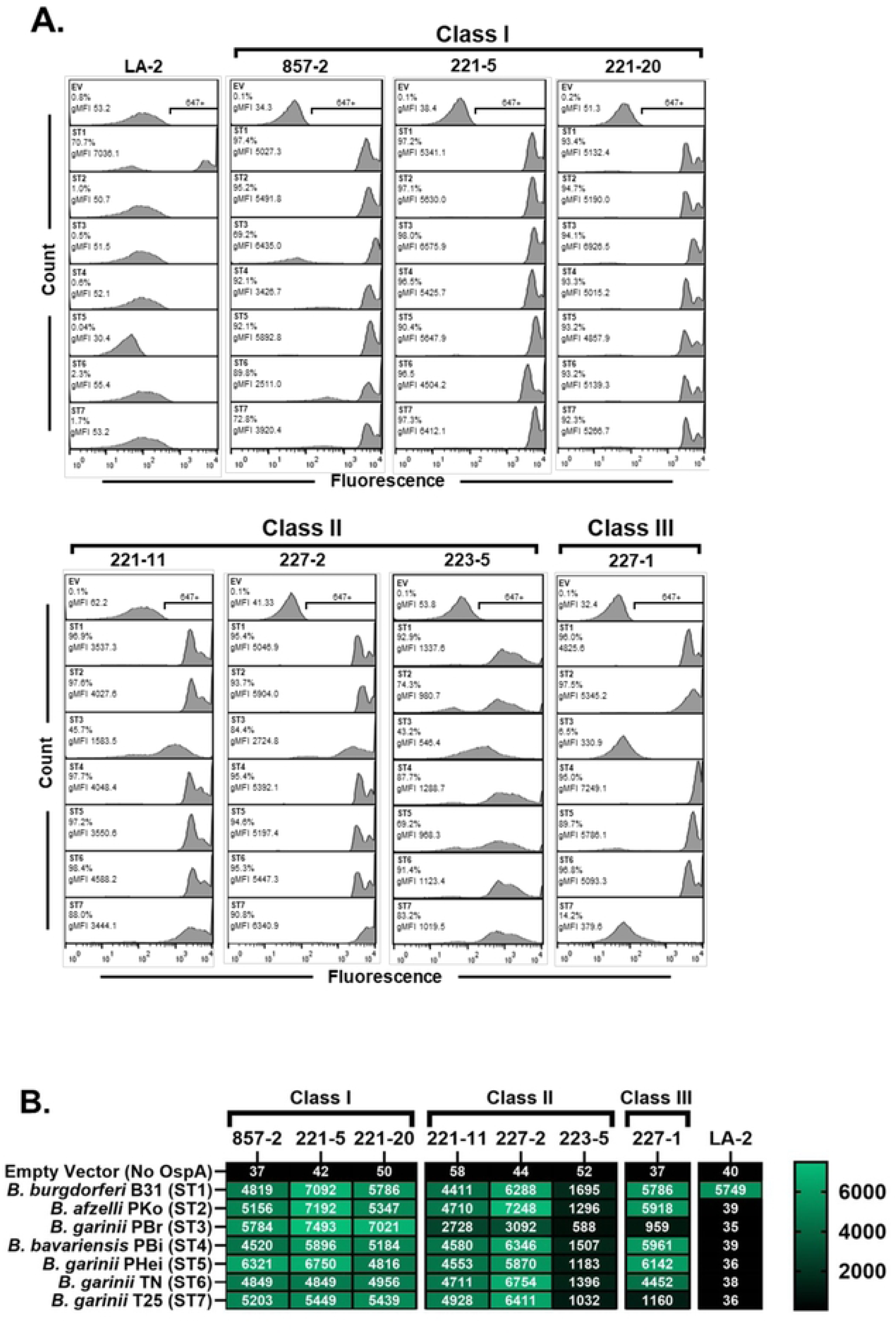
Surface binding profiles of anti-OspA_ST1_ Bin 1 mAbs to *B. burgdorferi* OspA ST1-7. (**A**) Representative flow cytometric histograms of live *B. burgdorferi* HB19-R1 reporter strains expressing OspA ST 1-7, each probed with 10 µg/ml of mAbs indicated in bold text above each column, followed by Alexa 647-labeled goat anti-human IgG secondary antibody. HB19-R1 harboring the empty vector (EV) was used as a negative control. The horizontal bracket represents the region of events positive for fluorescence labeling (647+) on the subsequent plots. The percent (%) of events positive for Alexa 647 fluorescence labeling and the geometric mean fluorescence intensity (gMFI) are shown in the top left corner of each box. Panels and plots were assembled using FlowJo. (**B**) Heat map summarizing the average gMFIs (with scale shown on right) of *B. burgdorferi* HB19-R1 reporter strains probed with each of the different mAbs depicted in Panel A. The results per box are the average of three biological replicates.

The HB19-R1 reporter strains were then probed with the additional Bin 1 mAbs in our collection to assess breadth of OspA serotype recognition (**Table 1**). Interestingly, the mAbs fell into three distinct classes (**Figure 1**). Class I mAbs (857-2, 221-5, 221-20) reacted strongly with OspA_ST1–7_ with MFIs >4000, while the Class II mAbs (221-11, 227-2, 223-5) had diminished recognition of OspA_ST3_. A single mAb, 227-1, was defined as Class III because of significantly reduced binding to OspA_ST3_ and OspA_ST7_. To determine if these class designations are a function of mAb binding to the different OspA serotypes and not an artefact of expressing heterologous OspA types on the surface of HB19-R1, we evaluated mAb binding to recombinant OspA_ST1–7_ by microsphere immunoassays (MIA). The mAb binding profiles to recombinant OspA_ST1–7_ mirrored exactly those observed with the HB19-R1 reporter strains (**Figure S2**). These results indicate that a subset of Bin 1 mAbs, namely those designated as Class I (857-2, 221-5 and 221-20), recognize epitopes that are conserved across the major OspA serotypes.

**Table 1.** Characteristics of OspA mAbs used in this study.

| mAb | bin<sup>a</sup> | class<sup>b</sup> | K<sub>D</sub><sup>c</sup> | iGL<sup>d</sup> V<sub>H</sub> | iGL V<sub>L</sub> | HCDR3 | PDB |
| --- | --- | --- | --- | --- | --- | --- | --- |
| 221-7<sup>e</sup> | 1 | (2) | 0.66 | HV5-51 | KV1-13 | ARGILRYFDWFLDY | 7JWG |
| 857-2 | 1 | 1 | 0.30 | HV5-51 | KV1-13 | ARHITTHTYRGFFDF | 8TUZ |
| 221-5 | 1 | 1 | 0.52 | HV5-51 | KV3-11 | ARHLGIDYFFGMDV | 8TV3 |
| 221-20 | 1 | 1 | 0.37 | HV5-51 | KV3-11 | ARHLGIDYFFGLDV | n.a. |
| 221-11 | 1 | 2 | 0.35 | HV5-51 | KV1-13 | VRGIGRYSNWFLDY | 8TVD |
| 227-2 | 1 | 2 | 0.48 | HV5-51 | KV1-13 | VRSISRSGPGYILSWFFDY | n.a. |
| 223-5 | 1 | 2 | 0.17 | HV5-51 | KV1-13 | ARYVTVGAFDF | n.a. |
| 227-1 | 1 | 3 | 1.7 | HV5-51 | KV1-13 | TRSISRSGPGFSLSWFFDY | 8TVJ |
| LA-2 | 3 | n.a. | 64 | n.a. | n.a. | n.a. | 1FJ1 |

<sup>a</sup>, from Haque et al 2022; <sup>b</sup>, n.a. not applicable, n.d. not determined; <sup>c</sup>, nM; <sup>d</sup>, inferred germline sequence; <sup>e</sup>, from Schiller et al, 2022; 221-7 inferred to be in class 2, based on Figure S2;

### Cross-serotype borreliacidal activity associated with Bin 1 mAbs

We next wished to determine the functional capacity of the Bin 1 mAbs to promote complement-dependent borreliacidal activity across the different OspA serotypes. To validate the HB19-R1 OspA reporter strains for this purpose, we compared complement-dependent killing of an infectious *B. burgdorferi* B31-5A4-based reporter strain (34) to the HB19-R1 OspA_ST1_ derivative. The assays were performed under conditions optimized for each strain, due to slight differences in growth kinetics and intrinsic complement sensitivities. Cultures were mixed with serial mAb dilutions (0.08-10 nM) in medium containing 10% (B31-5A4) or 2.5% (HB19-R1) guinea pig complement. The following day, IPTG was added to the cultures to induce *mScarlet-I* expression and MFI was measured 24 h (HB19-R1) or 48 h (B31-5A4) later, as described in the Materials and Methods. Under these conditions, the five Bin 1 mAbs had EC₅₀ values ranging from 0.63 to 1.25 nM against both the B31-5A4 and HB19-R1 OspA_ST1_ reporter, while an IgG isotype control (PB10) had no effect on spirochete viability (**Figure 2A-B**). Moreover, the EC_50_ values for 857-2 derived from the HB19-R1 OspA_ST1_ strain are virtually identical to those reported for *B. burgdorferi* B31 evaluated using a completely different viability assay (11). Thus, the HB19-R1 reporter strains served as reliable readouts of OspA-mediated complement-dependent borreliacidal activity.

**Fig. 2.**
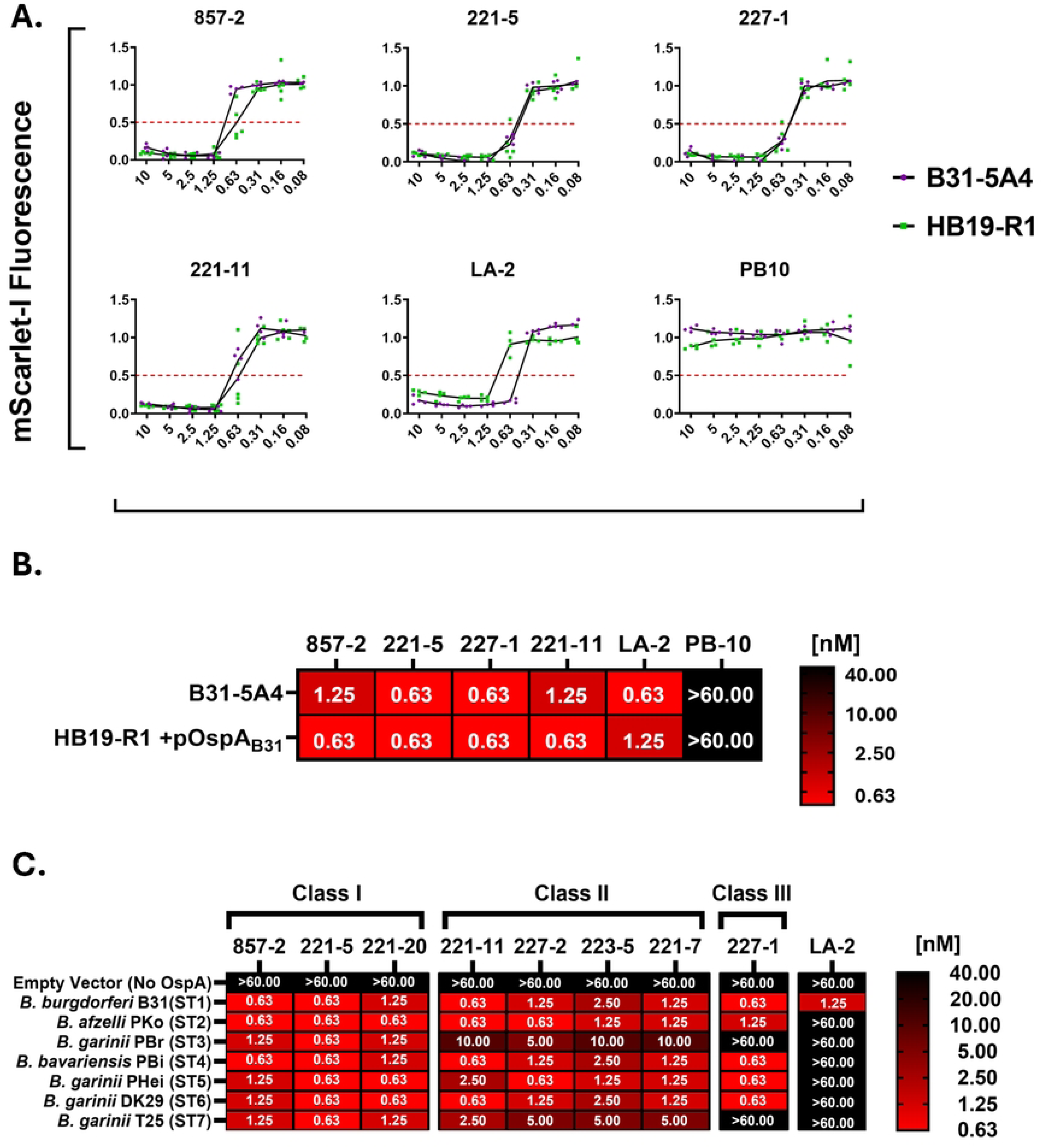
Anti-OspA_ST1_ Bin1 mAbs promote complement-dependent killing of recombinant *B. burgdorferi* Strains expressing OspA ST1-7. (A) Borreliacidal antibody titration curves and heat map **(B)** comparing the susceptibilities of an infectious *B. burgdorferi* B31-5A4 mScarlet-I viability reporter strain (34) and a recombinant HB19-R1 strain expressing the same OspA_ST1_ (B31) allele to complement-dependent killing by OspA_ST1_ mAbs. An irrelevant IgG1 (PB10), was also included as a control. Experiments were performed under conditions optimized for each strain as described within the Materials and Methods. Ec_50_ values shown represent the mean minimum concentration of antibody (nM) resulting in >50% reduction in MFI relative to controls. The data shown encompasses 3-5 independent experiments per strain with data normalized as described within the Materials and Methods section. **(C**) Heat map depicting the mean Ec_50_ (nM) of Bin1 mAbs against recombinant *B. burgdorferi* strains expressing OspA ST 1-7. Complement-dependent borreliacidal assays were performed with anti-OspA_ST1_ Bin1 mAbs and a panel of *B. burgdorferi* HB19-R1 (*ospA* negative) strains harboring an IPTG-inducible mscarlet-I viability reporter plasmid and expressing *ospA* ST1-7 under conditions described in the materials and methods section. Controls included an HB19-R1 strain carrying the IPTG-inducible viability reporter plasmid alone (empty vector, *ospA* negative) and a mAb with borreliacidal activity restricted to OspA_ST1_ (LA-2). Ec_50_ values shown represent the mean lowest concentration of antibody (nM) resulting in >50% reduction in MFI relative to controls. Reporter strains that exhibited resistance to complement-mediated killing by mAbs at 10 nM were later retested at a single higher dose (66.6 nM). The data shown encompasses 3-5 independent experiments per strain with data normalized as described within the materials and methods section. Antibody titration curves summarizing borreliacidal activity against each HB19-R1 reporter strain are shown in **Fig.S3.**

We therefore used the panel of HB19-R1 OspA reporters to assess the ability of the full collection Bin 1 mAbs (8 in total) to confer cross-serotype, complement-dependent borreliacidal activity (**Figure 2C**). The results reflected the OspA binding profiles in that **Class I** mAbs (857-2, 221-20, and 221-5) exhibited potent borreliacidal activity against all seven OspA serotypes, with EC₅₀ values ranging from 0.63 to 1.25 nM. In fact, 857-2’s EC_50_ values against the ST4 reporter strain match those reported for *B. bavariensis* strain (PBi) from which the OspA_ST4_ sequence was derived (11). **Class II** mAbs (221-11, 223-5, 227-2, and 221-7) had reduced potency against OspA ST3 and ST7 (EC_50_ 2.5-10 nM), while 227-1 (**Class III**) was completely devoid of ST3 and ST7 killing.

### Structural basis of OspA_ST1_ recognition by Bin 1 mAbs

To elucidate the epitopes recognized by the three different classes of Bin 1 mAbs, we solved the X-ray crystal structures of FAbs 857-2 (**Class 1**), 221-5 (**Class 1**), 221-11 (**Class 2**), and 227-1 (**Class 3**) in 1:1 stoichiometric complexes with OspA_ST1_ at resolutions ranging from 2.2 Å to 3.2 Å (**Table 2**; **Figure 3; Table S3**). After molecular replacement, the resulting phase information was used to calculate electron density maps employed to manually insert the corresponding residues into each model while manually building additional regions within each Fab-OspA model. Refinement of the four Fab-OspA structures revealed molecular models that possessed excellent geometry and that were consistent with the crystallographic data (**Table S4**).

**Fig. 3.**
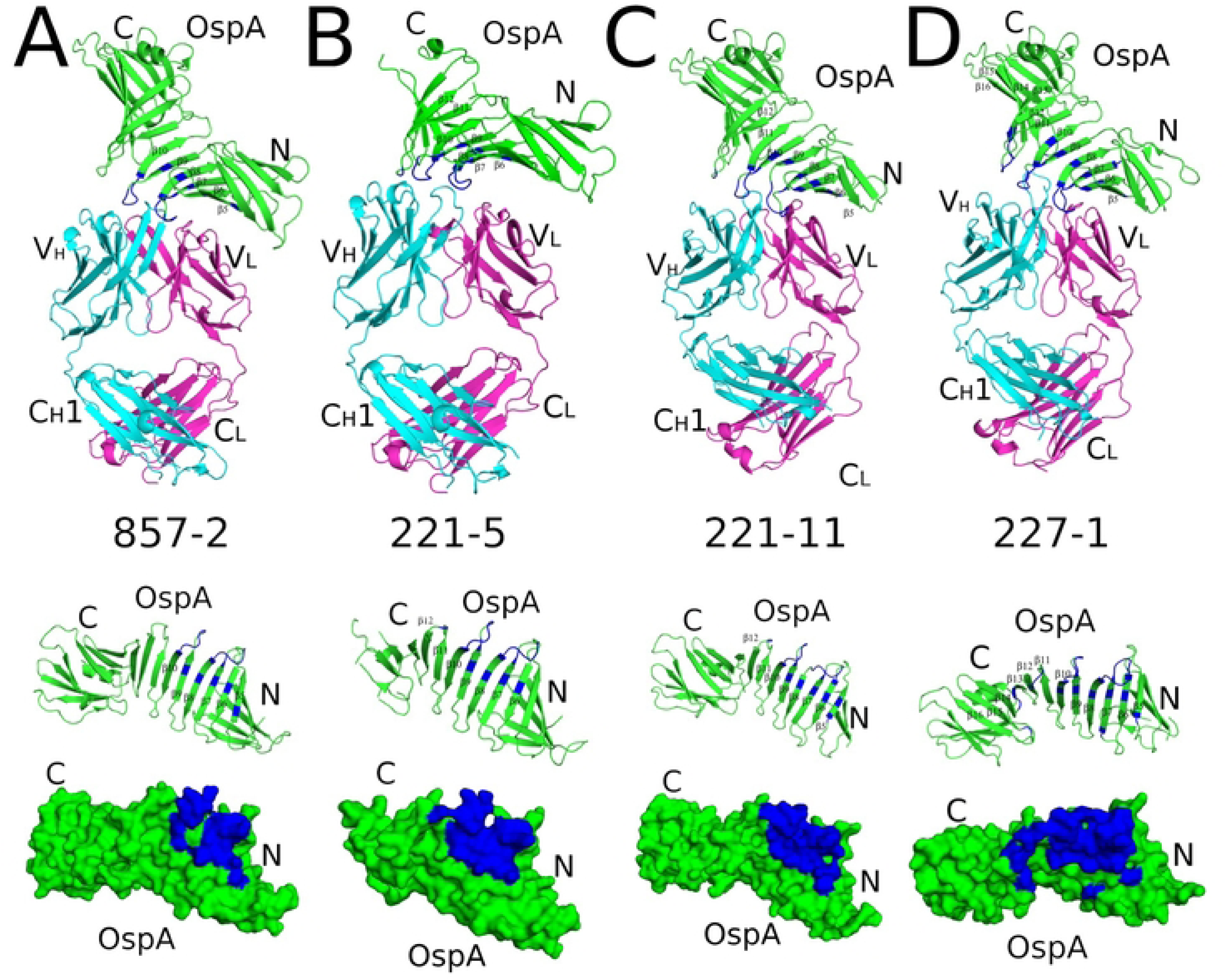
Crystral structures of Fab-OspA complexes reveal conserved epitopes on OspA_ST1_. (**top row**) Ribbon diagrams of OspA_ST1_ (green) in complex with Fabs from (A) 857-2, (B) 221-5, (C) 221-11 and (D) 227-1. Fab heavy chains (V_H_ and C_H_1) are colored cyan and light chains (V_L_ and C_L_) are colored magenta. The OspA N- and C-termini are labelled accordingly, along with selected strand numbers. (**middle row**) The four OspA (green) ribbon representations aligned in the same orientation with respective Fab contract points colored in blue. (**bottom row**) The ribbon and surface representations of OspA highlight the Fab-interacting residues, shown in blue. The N and C-termini of OspA are labelled N and C, respectively.

**Table 2.**
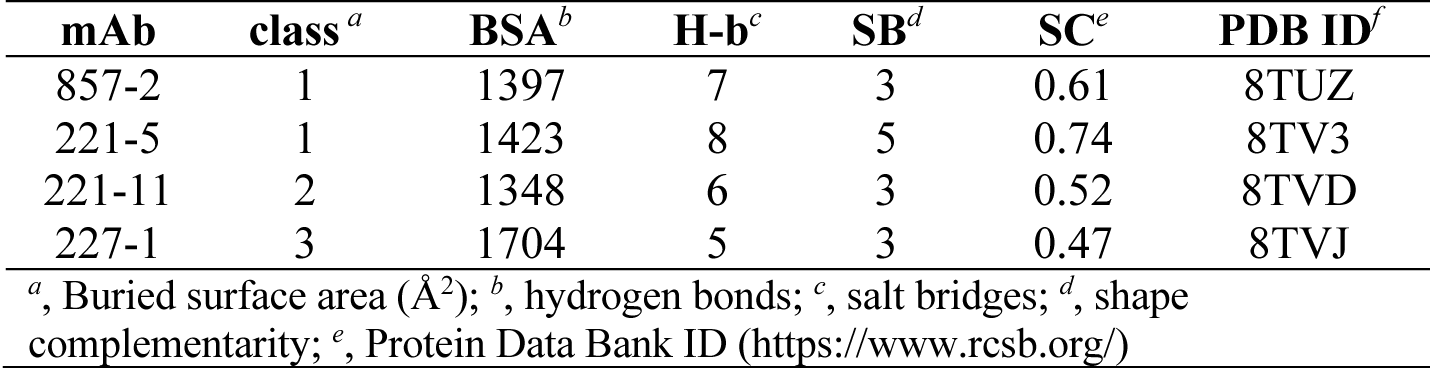
Summary of OspA-Fab interactions.

In each of the four complexes, the Fabs assumed a canonical structure with two heavy chain immunoglobulin domains (V_H_, C_H_1) and two light immunoglobulin domains (V_L_, C_L_) each containing 7-9 β-strands arranged in two β-sheets that folded into a two-layer sandwich with all six CDRs (L1-3; H1-3) on one face of each molecule. OspA was comprised of one antiparallel β-sheet with 21 β-strands (referred to as β-strands 1 to 21) connecting globular N- and C-terminal domains with a single α-helix (referred to as α-helix A) at the C-terminus (**Figure 3**). All 21 β-strands were present in their respective electron density maps of OspA complexed with Fabs 227-1 and 857-2. In the OspA-221-11 complex β-strands 1 and 2 were not visible in the electron density maps, while β-strands 17-21 were absent in the electron density maps of the OspA-221-5 complex, likely because of greater flexibility within these regions of OspA when bound to the particular Fabs. The OspA molecules from each of the four complexes were structurally similar to OspA alone (PDB ID: 2G8C), as evidenced by Root Mean Square Deviation (RMSD) of 0.8 Å to 2.4 Å upon superposition (**Figure S3**). Thus, the Fabs did not induce any major conformational changes within OspA.

The OspA-Fab structures revealed that 857-2 (**Class I**), 221-5 **(Class I**), 221-11 (**Class II**), and 227-1 (**Class III**) recognize overlapping but distinct epitopes on OspA with similar modes of engagement (**Figure 3**). The four Fabs contact OspA β-strands 6-10 and loop regions between β-strands 7-8 (loop 7-8) and 9-10 (loop 9-10), including residues 87, 100, 102, 105, 106, 107-109, 112, 124, 126, 128-132 (**Figure 3**). All but 857-2 Fab contacts OspA loop 11-12.

Fab 857-2 (**Class I;** K_D_ 0.30 nM) and 221-5 (**class 1;** K_D_ 0.52 nM) buried 1,397 Å² and 1,423 Å^2^, respectively, involving CDRs H1, H2, H3, and L2. 857-2 formed seven hydrogen bonds and three salt bridges with OspA, while 221-5 forms 8 hydrogen bonds and five salt bridges with OspA (**Table 2**). The Fabs each form two salt bridges with OspA residue Lys-107; one via CDR-L2 residue Asp/Glu-55 and the other with CDR-H3 residue Asp-109/110 (**Figure 4A-B**). Fab 221-5 makes contacts with Ser-152 in loop 11-12 via framework region (FR) residue Arg-59 but is too distant (4.2 Å) to form an H-bond. 857-2 does not interact with loop 11-12 at all, as Arg-59 is >9 Å from OspA Ser-152.

**Fig. 4.**
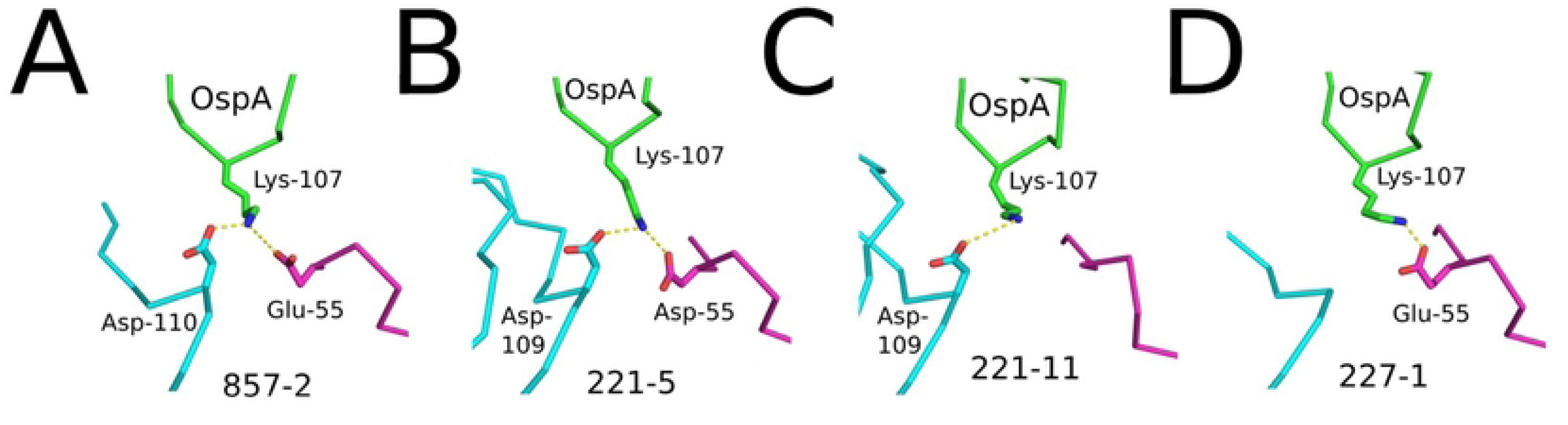
Key interactions between each Fab and OspA. Close-up of notable salt bridges between OspA Lys-107 (green) and Fab residues are shown as yellow dashed lines. Fab heavy chains (V_H_/C_H_1) are colored cyan, and light chains (V_L_/C_L_) are colored magenta. (A) Fab 857-2 engages Lys-107 through Glu-55 and Asp-110. (B) Fab 221-5 interacts with Lys-107 via Asp-55 and Asp-109. (C) Fab 221-11 contacts Lys-107 through Asp-109. (D) Fab 227-1 engages Lys-107 through Glu-55. All side chains are drawn as sticks and color coordinated to the main chain color with nitrogen atoms blue and oxygen atoms red.

Fab 221-11 (**Class II;** K_D_ 0.35 nM) engages with OspA via CDRs H1, H2, H3, and L2 and has a total buried surface area of 1,348 Å^2^ (**Table 2**). It forms 9 polar contacts with OspA, including a hydrogen bond between CDR-H1 Tyr-32 and OspA Asp-105. Like 857-2 and 221-5, 221-11 also forms a salt bridge between CDR-H3 residue Asp-109 and OspA residue Lys-107 (**Figure 4C**). However, unlike the other two mAbs, 221-11 light chain FR Glu-55 does not form a second H-bond with Lys-107 (**Figure 4C**).

Fab 227-1 (**Class III;** K_D_ 1.7 nM) stands out from the other three Fabs in that it forms the most extensive interface with OspA, burying a total of 1,704 Å² with contributions from five of the six CDRs (H1-3, L1-2). Fab 227-1 establishes five hydrogen bonds and three salt bridges with OspA, including with Lys-107 (**Figure 4D**). FR residue Arg-59 forms a hydrogen bond with Ser-152 of OspA, as will be discussed in detail below. 227-1 makes additional contacts with OspA not observed in the other three Fabs, including with loop 11-12 (residues 153-154), β-strand 13, loop 13-14 (residues 171-173), and loop 15-16 (residue 193). These additional interfaces result in 239 Å² of total buried surface area. This larger binding surface likely comes at a cost, considering that 227-1 (**Class III**) has a restricted OspA serotype profile (i.e., fails to bind OspA_ST3_ and OspA_ST7_) relative to 857-2 and 221-5, which bind the seven major OspA types.

### Structural basis of cross serotype restriction of Bin 1 mAbs

Based on the structural analysis of the OspA-Fab interactions, we postulate that two distinct molecular interactions account for the reduced binding of 221-11 (**Class II**) and 227-1 (**Class III**) to OspA ST3 and ST7. The first interaction involves the Glu_ST1_>Lys_ST3/7_ polymorphism at position 131 (**Figure 5A, B)**. For example, 227-1 V_H_ FR3 residue Arg-59 forms a salt bridge with Glu-131 in OspA_ST1_, which would be disrupted by the substitution of a Lys at that position. Moreover, a mutation to Lys at position 131 would introduce electrostatic repulsion with Lys-129, which is located ∼5 Å away. As a result, the salt bridge between OspA Lys-129 and 227-1 V_H_ residue Asp-55 would be perturbed (not shown). Thus, a single Glu_131_Lys polymorphism would result in the loss of two salt bridges in the case of 221-11 and 227-1. Second, the Ser_ST1_>Asn_ST3/7_ polymorphism at position 152 may restrict 221-11 (**Class II**) and 227-1 (**Class III**) recognition of OspA ST3 and 7. 221-11 and 227-1 each form H-bonds with OspA_ST1_ residue Ser-152 via V_H_ residue Arg-59 (**Figure 5C,D**). The Asn polymorphism at this position would perturb a corresponding interaction in OspA_ST3_ and OspA_ST7_.

**Fig. 5.**
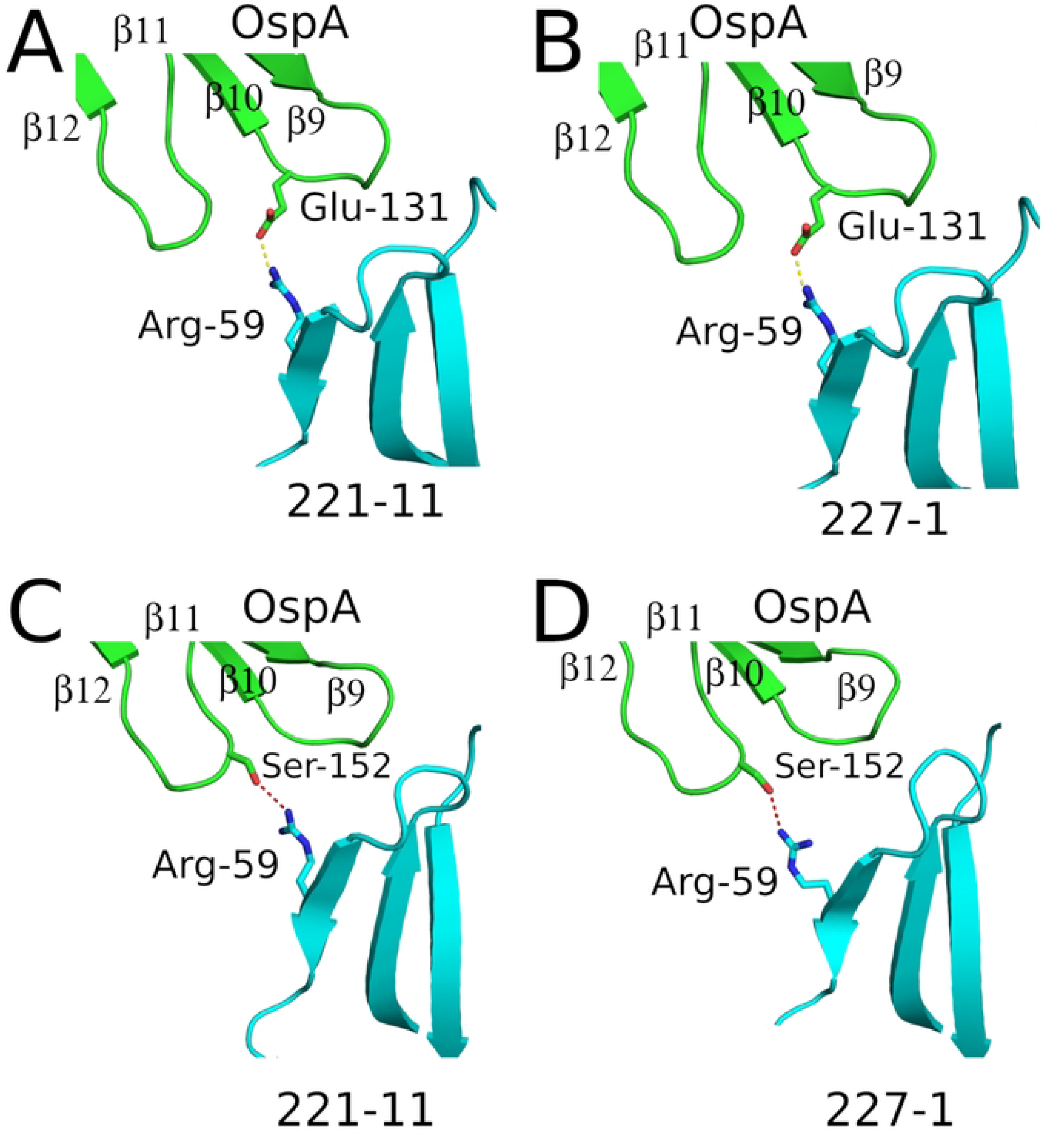
Influence of OspA primary sequence deviations on antibody function. (A) Structural basis for Fabs 221-11 and 227-1 cross-reactivity loss in OspA serotypes 3 and 7.Distinct hydrogen bond networks between Fab Arg-59 (cyan) and OspA residues (green) contribute to serotype specificity. (A) In Fab 221-11, Arg-59 forms a salt bridge with OspA Glu-131. (B) The same Glu-131 interaction is observed in Fab 227-1. (C) In 221-11, Arg-59 hydrogen bonds with OspA Ser-152. (D) Similarly, Fab 227-1 also engages Ser-152 through Arg-59. The primary sequence difference distinguishing serotypes (STs) 1, 2, 4, 5, and 6 from STs 3 and 7 involves a Glu-Lys substitution at position 131 and a Ser-Asn at position 152. These two primary sequence differences conceivably disrupt the Arg-59–Glu-131 salt bridge and the Arg-59 hydrogen bond with Ser-152, consequently reducing the cross-reactivity of Fabs 221-11 and 227-1 with ST3 and ST7. All side chains are drawn as sticks and color coordinated to the main chain color with nitrogen atoms blue and oxygen atoms red. Salt bridges are represented as yellow dashes and hydrogen bonds are red dashes.

### OspA recognition is driven by V_H_ and V_L_ germline-encoded residues

All eight of the Bin 1 mAbs, including the four whose structures were solved in complex with OspA (857-2, 221-5, 221-11, 227-1) as well as the previously reported 221-7 (PDB 7JWG), are inferred to utilize the same HV5-51 germline lineage (**Table 1**; **Figure S6**). In the case of 221-5 (**Class I**), the V_H_ is unmutated from the HV5-51 germline, while the V_H_ elements of the other four mAbs with solved co-complexes have only nominal mutations (4-6 amino acids) that are not associated with direct OspA interactions. Additionally, the H-CDR3 residue involved in the well-conserved salt bridge with Lys-107, namely Asp109 (or Asp110 or 114, depending on CDR3 length), is encoded by the J_H_ gene segment in five of the six human germline J genes (J_H_1 being the exception) (**Table 1**). Similarly, in six of the mAbs, the V_L_ is derived from the KV1-13 with germline residues involved in critical contacts, including Glu-55 that forms a salt bridge with OspA Lys-107 (**Figure 4**). In the case of 221-5 and 221-20 (**Class I**), which are derived from KV3-11, a single mutation in codon 55 (C>A, resulting in amino acid mutation Ala>Asp) would be sufficient to form a salt bridge with OspA Lys-107. Therefore, BCR combinations with HV5-51 and either KV1-13 or KV3-11 would be expected to have broad affinity for OspA serotypes.

### Borreliacidal epitopes are conserved across OspA in diverse *Bbsl* genospecies

With a detailed structural understanding of the interactions of Bin 1 mAbs with OspA_ST1_, we sought to examine epitope conservation across the *Bbsl* genospecies complex. To accomplish this, we surveyed OspA protein sequences throughout the *Boreliella* genus to identify novel OspA types beyond the eight STs and nine ISTs that have been reported (37). To this end, we constructed an unrooted phylogenetic tree consisting of 135 OspA protein sequences collected from all 23 recognized *Bbsl* genospecies (38). To facilitate the identification of clades corresponding to known ST/ISTs, we included 86 OspA reference sequences obtained from the PubMLST OspA IST collection assembled by Lee (37).

The resulting phylogenetic tree revealed 33 distinct sequence clusters corresponding to OspA ST 1-8, IST 9-17, and 16 other sequence groups representing previously untyped OspA variants (**Figure 6A**). The 86 OspA reference sequences grouped into discrete clades consistent with their previously designated ST or IST designations (37). The one exception was *B. valaisiana*, which clustered into two distinct clades: four sequences clustering as IST15, and three other sequences sharing close homology with IST16, principally associated with *B. turdi* (**Figure 6A**). We also identified a single *B. garinii* OspA sequence (isolate IPT75) that grouped independently of the eight recognized OspA types found within this genospecies. The 14 other OspA clades identified corresponded to untyped *B. burgdorferi* sl genospecies, including *B. americana, B. andersonii, B. bissettiae, B. californiensis, B. carolinensis, B. chilensis, B. finlandensis, B. japonica, B. kurtenbachii, B. lanei, B. lusitaniae, B. maritima, B. sinica,* and *B. tanuki* (**Figure 6A**).

**Fig. 6.**
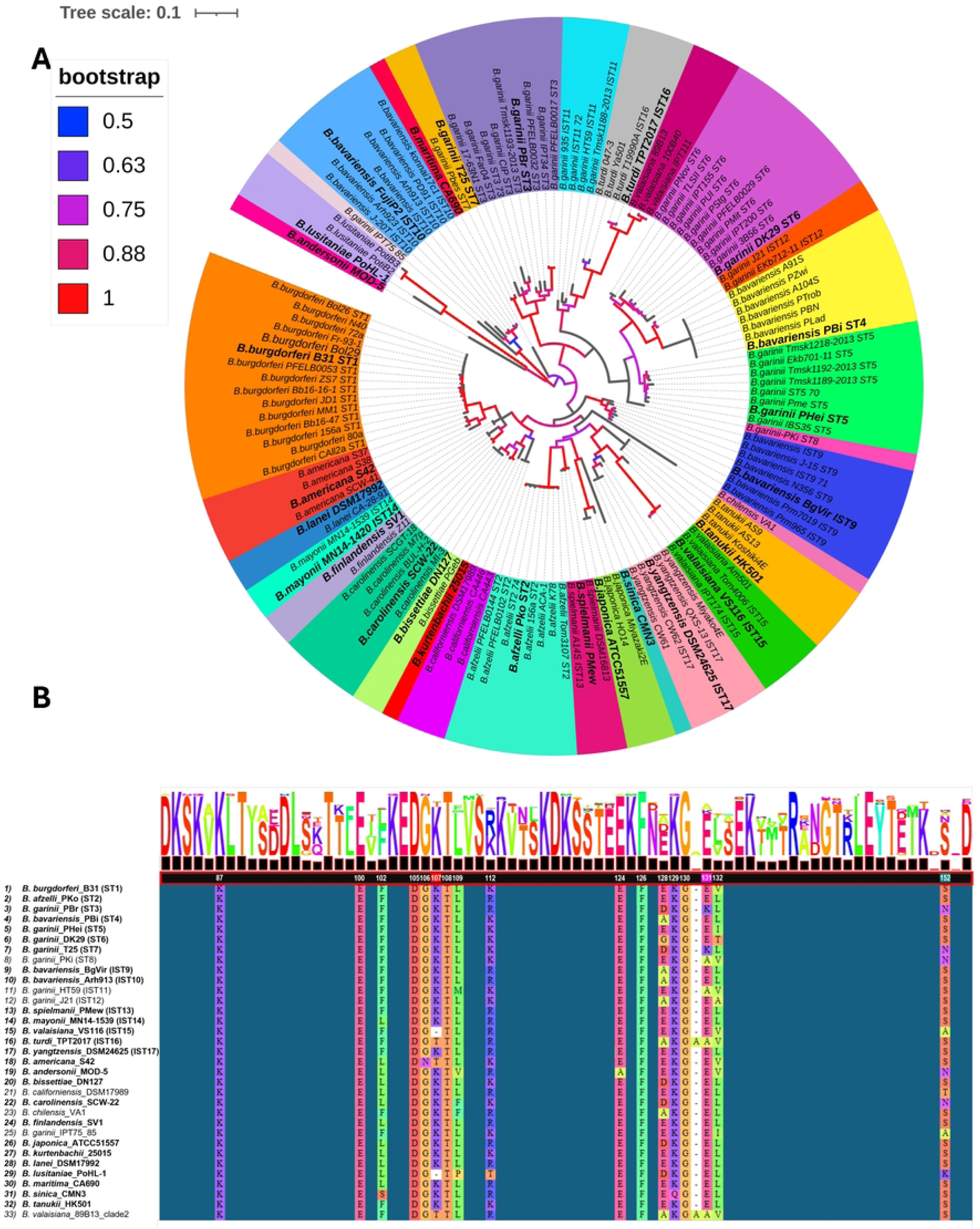
Phylogenic analysis of OspA within the *Borreliella burgdorferi sensu lato* genospecies complex. **(A)** Unrooted phylogenetic tree comprised of 135 OspA protein sequences from 23 *Bbsl* genospecies. Phylogenetic analysis identified 33 distinct OspA clades corresponding to serotypes (ST) 1-8, *in silico* types (IST) 9-17, and 16 other unclassified OspA IST’s. Clades representing each distinct OspA variant/type are distinguished by color. Enlarged and bolded strain names denote OspA alleles selected for HB19-R1 viability reporter strain construction. Bootstrap values representing approximately maximum likelihood analysis are distinguished by color: >0.50 = green, >0.75 = blue, 1.00 = red. Nodes with bootstrap values under 0.50 are not shown (gray) **(B)** Simplified multiple sequence alignment of the Bin1 epitope region of 33 unique OspA types. Representative sequences from each OspA clade were aligned using OspA from *B. burgdorferi* strain B31 as a reference. Bolded strain names indicate OspA variants expressed by HB19-R1 viability reporter strains. Amino acid residues numbered in white font denote conservation within two or more epitopes of Bin1 mAbs 857-2, 221-5, 221-11, 221-7, and 227-1. Numbered amino acid residues with colored background form critical interactions with residues within the paratopes of Bin1 mAbs with solved crystal structures (Lys-107, Glu-31, Ser-152).

### Genetically diverse OspA types are susceptible to Bin 1 mAbs

To assess epitope conservation across the *Bbsl* genospecies complex, we generated a multiple sequence alignment (MSA) of representative OspA sequences from each of the 33 clades identified in our phylogenetic analysis. Using OspA ST1 as the reference, the alignment revealed broad conservation of the loop regions between β-strands 7-8 and 9-10, including residues associated with 857-2 and other Bin1 mAb epitopes (**Fig. 6B; Fig. S7**).

To experimentally examine conservation of 857-2’s and other Bin 1 epitopes beyond OspA ST 1-7, we generated 19 additional HB19-R1 reporter strains expressing OspA variants from two *B. bavarienisis* types (IST9, IST10) and 17 other *Bb*sl genospecies. Many of the OspA types selected are occasionally associated with Lyme disease in humans such as *B. bavariensis* IST9 and IST10, *B. spielmanii* (IST13), *B. mayonii* (IST14), *B. americana*, *B. bissettiae*, and *B. lusitaniae* or exhibit limited or uncertain disease potential in humans like *B. andersonii,* and *B. yangtzensis* (38–41). We also generated HB19-R1 viability reporter strains expressing OspA types from non-pathogenic Bbsl genospecies to fully capture Bbsl complex-wide sequence diversity and explore the impact of polymorphisms predicted to disrupt Bin 1 mAb binding.

These included *B. valaisiana* (IST15), *B. turdi* (IST16), *B. sinica*, *B. finlandensis*, *B. japonica*, *B. lanei*, *B. carolinensis*, *B. kurtenbachii*, *B. maritima*, and *B. tanuki* (38, 40, 42). The source strains for each OspA type or IST are indicated in bold within the phylogenetic tree (**Figure 6A**) and corresponding multiple sequence alignment (**Figure 6B, S7).**

The Bin 1 mAbs were then evaluated for complement-dependent borreliacidal activity against the 19 additional OspA reporter strains described above. **Class I** mAbs (221-20, 221-5, 857-2) were the most broadly functional, as exemplified by 221-20, which exhibited potent borreliacidal activity against 23 of the 26 OspA reporter strains within our collection [EC_50_ 1.25-2.50 nM] (**Figure 7**). The three resistant strains (EC_50_ >60 nM) were *B. turdi* (IST16), *B. lusitaniae,* and *B. sinica*. 221-5 and 857-2 susceptibility profiles were similar to 221-20, with the addition of both having lessened activity against *B. valaisiana* (IST15), and 857-2 also being ineffective against *B. americana* (**Figure 7**). The **Class II** mAbs (221-11, 227-2 and 223-5) had borreliacidal activity profiles similar to 857-2, albeit with reduced potencies (**Figure 7**).

**Fig. 7.**
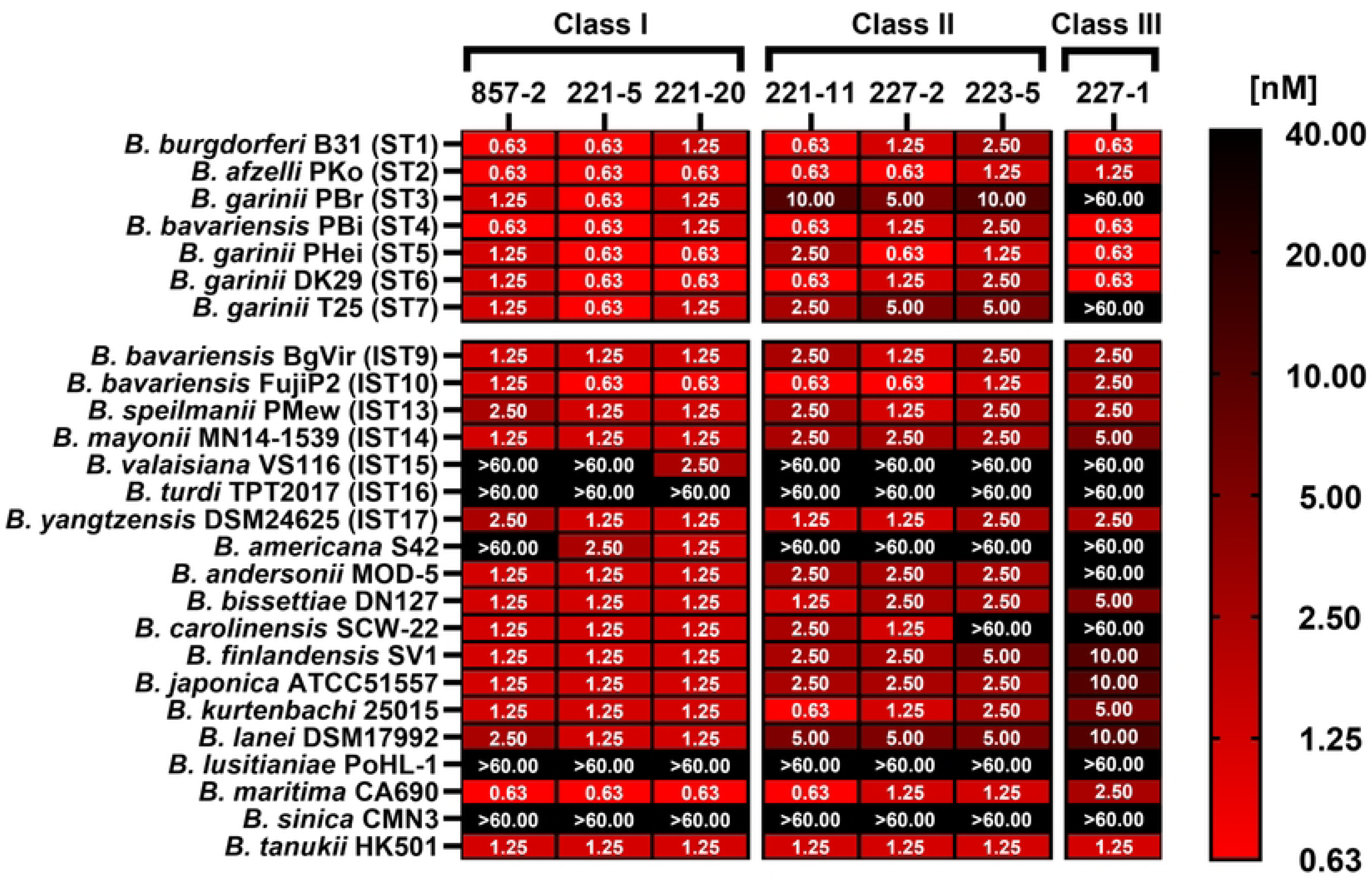
Genetically diverse OspA types are susceptible to complement-mediated killing by anti-OspA_ST1_ Bin1 mAbs. Complement-dependent killing assays were performed as described in the materials in methods section with anti-OspA Bin1 mAbs and a panel of *B. burgdorferi* HB19-R1 viability reporter strains expressing 19 OspA *in silico* types (ISTs). (A) Heat map summarizing the mean EC_50_ values for complement-mediated killing by Bin1 mAbs from classes 1–3. The susceptibility profiles of reporter strains expressing OspA ST1–7 are shown for comparison. Reporter strains that exhibited resistance to complement-mediated killing by mAbs at 10 nM were later retested at a single higher dose (66.6 nM). EC₅₀ values denote the lowest antibody concentration (nM) capable of reducing mScarlet-I fluorescence by >50% relative to untreated controls and represent the mean of at least three independent experiments. Antibody titration curves summarizing borreliacidal activity against each strain are provided in **Fig.S8.**

Interestingly, 223-5 was distinct among this class for its inability to kill *B. carolinensis*. Finally, 227-1 (**Class III**) had borreliacidal activity against just 17 of the 26 OspA reporter strains and reduced efficacy (Ec_50_ 5-10 nM) against OspA variants from *B. mayonii* (IST14), *B. finlandensis, B. japonica,* and *B. lanei* (**Figure 7**). These results demonstrate that one or more borreliacidal epitopes are conserved across virtually all the major variants within the *Bbsl* genospecies complex.

### Lys-107 polymorphism as a determinant of susceptibility to nearly all Bin 1 mAbs

As noted above, HB19-R1 strains expressing OspA from *B. turdi* (IST16) and *B. lusitaniae* PoHL-1 were resistant to all the Bin 1 mAbs tested, while *B. americana* and *B. valaisiana* (IST15) were resistant to a subset of **Class I** mAbs and all the **Class II** and **Class III** mAbs (**Figure 7**).

Examination of the OspA MSA (**Figure 6B**) suggests that polymorphisms at OspA_ST1_ Lys-107 may account for all or part of the observed resistance. In the case of *B. turdi* (IST16), for example, there is a Thr residue rather than a Lys at position 107, while OspA from *B. lusitaniae* has a deletion at the equivalent position (**Figure 6B**). Polymorphisms at Lys-107 would preclude 857-2 and other mAbs from forming critical salt bridges with OspA (**Figure 4**).

To investigate the importance of Lys-107 in the context of OspA recognition by the Bin 1 mAbs, we generated HB19-R1 reporter strains expressing OspA_ST1_ variants harboring a K107T substitution or Lys-107 deletion (ΔK107). The OspA_ST1_ K107T and ΔK107 reporter strains were completely resistant to killing by the **Class II** (227-2, 223-5, 221-11) and **Class III** mAbs (227-1), even at the highest antibody concentrations tested (66.6 nM) (**Figure 8**). Similarly, both modifications were detrimental to **Class I** mAbs 857-2 and 221-5’s borreliacidal activity, as evidenced by 16-fold increases in EC_50_ values (**Figure 8**). For 857-2, the OspA_ST1_ ΔK107 deletion was slightly more deleterious than the K107T substitution (EC_50_ 20 nM vs 5 nM), while 221-5’s resistance was comparable between the two (10 nM). Conversely, the OspA_ST1_ K107T and ΔK107 reporter strains remained sensitive to killing by 221-20, demonstrating the unique nature of 221-20’s paratope-epitope interaction and differentiating 221-20 from the other two Bin 1 mAbs.

**Fig. 8.**
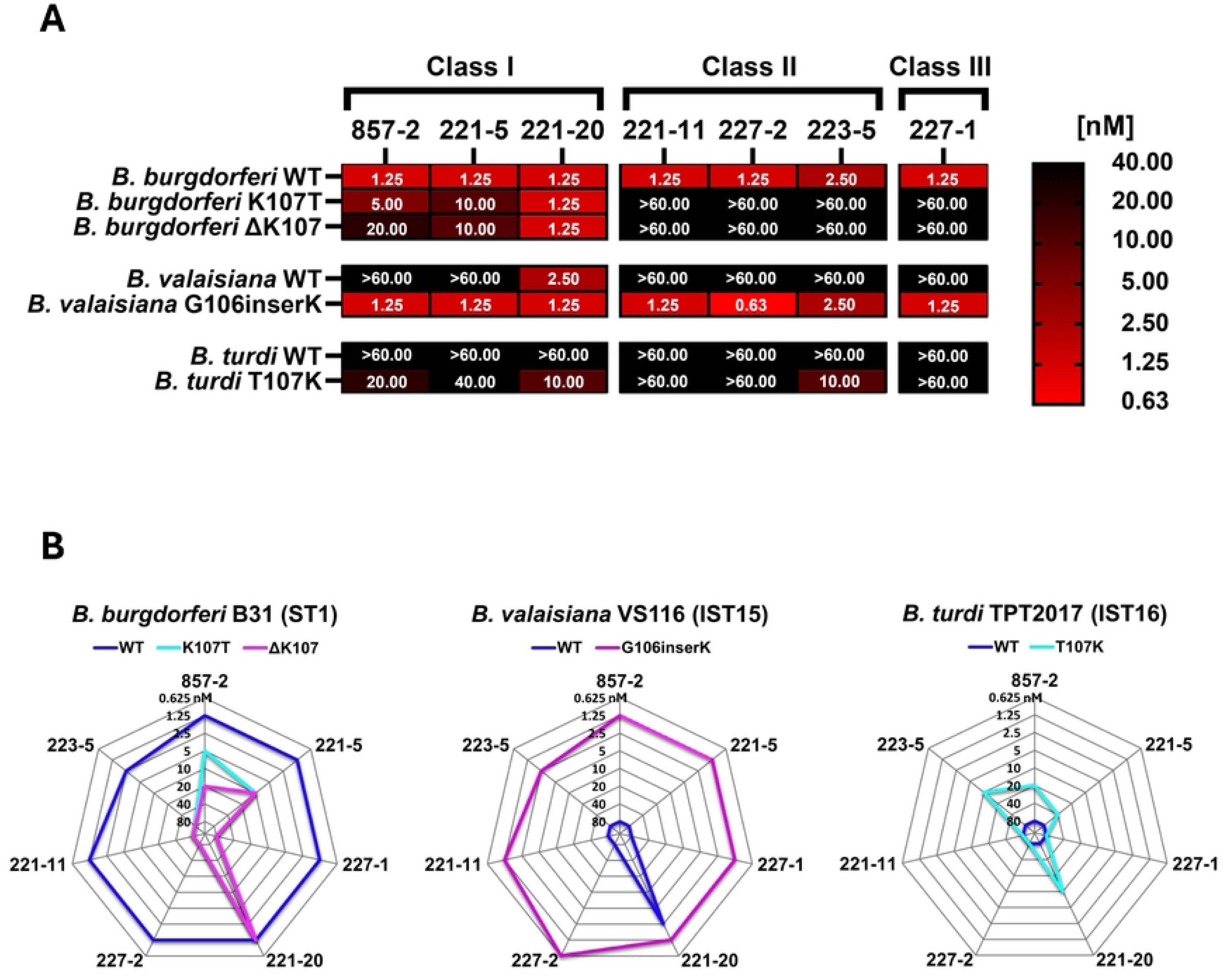
Lys-107 polymorphisms are a determinant of susceptibility to nearly all Bin1 mAbs. Complement-dependent killing assays were performed as described in the materials and methods section with anti-OspA Bin1 mAbs and *B. burgdorferi* HB19-R1 viability reporter strains expressing “WT” OspA variants from *B. burgdorferi* B31 (ST1), *B. valaisiana* VS116 (IST15), and *B. turdi* TPT2017 (IST16), or mutated derivatives containing deletions, insertions, or substitutions at residue 107. Heat maps (A) and radar plots (B) depicting the EC₅₀ profiles of WT and muta**t**ed OspA variants. EC₅₀ values denote the lowest antibody concentration (nM) capable of reducing mScarlet-I fluorescence by >50% relative to untreated controls and represent the mean of at least three independent experiments. Reporter strains exhibiting resistance to killing by mAbs at 20 nM were re-examined at 40 nM and 66.6 nM doses. Antibody titration curves summarizing borreliacidal activity against each strain are shown in **Fig.S9.**

In light of these results, we examined whether introduction of Lys-107 in an otherwise resistant OspA variant from *B. valaisiana* (IST15) was sufficient to render cells susceptible to killing by the Bin 1 mAbs. In accordance with that hypothesis, the OspA_IST15__G106insK reporter strain was susceptible to all Bin 1 mAbs with EC_50_ values comparable to OspA_ST1_ (Ec_50_ range 0.63-2.50 nM), thereby demonstrating that antibody recognition is transformed by the addition of a single Lys residue at position 107.

We also generated a reporter strain with a *B. turdi* OspA (IST16) allele encoding a T107K substitution to investigate the role of Lys-107 in the context of polymorphisms elsewhere in 857-2 and other Bin 1 mAb epitopes (e.g., residues 128-132). In fact, the T107K substitution was sufficient to render *B. turdi* OspA at least partially susceptible to the **Class I** mAbs (857-2, 221-5, 221-20) and one **Class II** mAb (223-5) with Ec_50_ values ranging from 10-40 nM (**Figure 8**). Collectively, these experiments demonstrate that Lys-107 is a key susceptibility determinant for virtually all Bin 1 mAbs.

## DISCUSSION

The development of cross-protective vaccines for Lyme disease has been challenging due to the antigenic variability of OspA among the major pathogenic *B. burgdorferi* genospecies in North America, Europe, and Asia. Efforts to identify conserved epitopes on OspA have been limited, despite evidence in the literature for their existence (17, 43). Wilske and colleagues, for example, described a mouse mAb (L31 1F11) capable of reacting with all seven OspA STs and more than 100 primary isolates (15). L31 1F11’s epitope was localized through a series of OspA truncations and Western blotting to residues 102-111, which corresponds to OspA’s central β-sheet (15). Twenty years later, Wang and colleagues “rediscovered” this region of OspA in their effort to identify broadly reactive human mAbs with the potential to serve as preexposure prophylactics for Lyme disease (11). Two of the mAbs, 857-2 and 221-7, were shown to promote complement-dependent killing of three different OspA serotypes—*B. burgdorferi* (ST1), *B. afzelii* (ST2) and *B. bavariensis* (ST4)—and passively protect mice and non-human primates (221-7) from *B. burgdorferi* (OspA_ST1_) tick-mediated challenge (11, 12). While 221-7’s epitope was reported by us several years ago (12), we reasoned that a more detailed and comprehensive analysis of this region of OspA would shed light on key epitope-paratope interactions associated with protection.

Indeed, in our current study we identified a protective B cell epitope shared across all the major OspA serotypes associated with Lyme disease-causing spirochetes. The core of the epitope involves residues within β-strands 6-10 and loops 7-8 and 9-10 (**Figure S10**). Within this region, there are invariant (e.g., Glu-100, Asp-105) and highly conserved residues that contribute to antibody recognition. Varying paratope interactions within and outside the core epitope likely account for the different degrees of mAb reactivity with the seven major OspA serotypes. We speculate, for example, that single amino acid polymorphisms relative to OspA_ST1_ likely account for the reduced binding of 221-11 (**Class II**) and 227-1 (**Class III**) to OspA_ST3_ and OspA_ST7_.

Conversely, 221-20 (**Class I**) stood out for its potent borreliacidal activity against the seven major OspA STs and 16 of the 19 OspA IST reporter strains tested. 221-20 was also the only mAb whose borreliacidal activity was unaffected by substitutions or deletions at OspA_ST1_ Lys-107. Unfortunately, efforts to solve the structure of 221-20 Fabs bound to OspA were unsuccessful, so the exact nature of 221-20’s paratope-epitope interaction remain undefined.

Our results suggest a predilection of particular V_H_ and VL combinations for recognition of OspA’s central β-sheet. Namely, the seven mAbs characterized in this study as well as 221-7 are all predicted to utilize V_H_ germline HV5-51 with critical epitope interactions mediated largely if not exclusively by germline encoded residues. Similarly, critical light chain contacts are associated with KV1-13 or KV3-11 germline residues, suggesting that particular BCR pairings (i.e., HV5-51 / KV1-13) may be sufficient to engage with conserved residues within OspA’s central β-sheet, including Lys-107. However, as these Bin 1 mAbs in this study were generated from hybridomas derived from hyperimmunized transgenic mice containing human immunoglobulin genes (11), we can only speculate that such interactions are clinically relevant. BCR repertoire analysis and the role of unmutated, class switch B cells and particular germline in response to Lyme disease has garnered attention recently (44–46). Jiang and colleagues examined BCR usage and frequency of somatic hypermutation in memory B cell populations from erythema migrans skin lesions and noted a subpopulation of unmutated memory B cells (45). Blum and colleagues examined BCR repertoires in circulating plasmablasts and, perhaps coincidently identified HV5-51 as being one of just two V_H_ germlines overrepresented in Lyme disease patients, as compared to healthy controls (44).

The generation of isogenic *B. burgdorferi* reporter strains expressing the seven major OspA serotypes (ST1–7) and numerous ISTs proved instrumental in being able to define cross-borreliacidal epitopes recognized by the seven mAbs in this study. We speculate that the same reporters would be equally valuable in assessing antibody responses elicited by OspA-based vaccines. At present, assessing complement-dependent bactericidal activity across different Bbsl genospecies requires the development of genospecies-specific assays in which culture conditions, complement sources, and assay duration must be optimized for each individual strain (47, 48). For example, because of *B. garinii*’s (ST3) inherent sensitivity to guinea pig, mouse and human complement, Comstedt and colleagues had resorted to freshly isolated chicken blood for bactericidal assays (25). Moreover, the overwhelming reliance on ATP-based viability assays is problematic due to the nature of *B. burgdorferi* metabolism and high signal to noise ratios. The adoption of a standardized fluorescence-based, serum borreliacidal assay (SBA) based on OspA reporter strains like those described herein would not only streamline the assay itself but would enable different manufacturers and regulatory agencies to compare vaccine efficacies using a single platform.

## MATERIALS AND METHODS

### Cloning and expression of OspA and anti-OspA mAbs

Recombinant human monoclonal IgG1 antibodies were purified via Protein A affinity column from Expi293 cells that had been transiently transfected with V_H_ and V_L_ plasmids, as described (11). Recombinant OspA_ST1_ (residues 18-273) from *B. burgdorferi* B31 (NCBI:txid224326) was expressed in *Escherichia coli* BL21-DE3 and purified as described (12). Purified OspA was maintained in 20 mM HEPES, 150 mM NaCl, 20 mM imidazole, pH 7.50. Recombinant OspA serotypes 1-7 purified from *E.coli* were kindly provided by Dr. Meredith Finn (Moderna) and described elsewhere (34).

### ELISA

Immulon 4HBX plates (ThermoFisher, Waltham, MA) were coated overnight at 4°C with 1 µg/mL of the seven OspA serotypes (ST1-7) and the next morning blocked for 2 h with 2% goat serum in PBS supplemented with 0.1% Tween-20 (PBST). Serial two-fold dilutions of each antibody starting from 1 µg/mL were made in separate dilution plates, and transferred to the serotype coated plates for 1 h. Plates were then washed three times with PBST, and a goat anti-human HRP conjugated secondary antibody (SouthernBiotech) was added for 1 h. Plates were then washed three times with PBST, developed with SureBlue TMB substrate (Seracare, Milford, MA), and quenched with 1M phosphoric acid. Absorption at 450 nm was read by a SpectraMax iD5 spectrophotometer using Softmax Pro v7.1 (Molecular Devices, San Jose, CA).

### Affinity measurements with BioLayer interferometry

Certain monoclonal antibody binding kinetics first reported by Haque and colleagues (32) have been revised to reflect refined and optimized BLI protocols using an Octet RED96e (Sartorius AG, Gottingen, Germany). AHC sensors (#18-5060, Sartorius) were used to capture mAbs at 0.4 µg/mL for 5 min, to a loading level of ∼0.5 nm, then dipped into wells containing a three-fold dilution series of OspA (ranging from 0.6 µg/mL, 21.58 nM to 0.0025 µg/mL, 0.089 nM) for 10 min. Finally, sensors were dipped into buffer alone for 30 minutes to measure dissociation. Buffer used in all wells was 0.1% BSA in PBS, with 0.01% Tween20. Sensors were regenerated before the first mAb and between each mAb with 0.2 M glycine, pH 2.2. Traces were background corrected with a reference sensor, in which mAb was loaded but not exposed to OspA, and a reference sample, in which a bare sensor was exposed to the highest concentration of OspA. Traces for each mAb were fit to a global 1:1 model. k_obs_ vs concentration was linear for each system.

### Construction of *B. burgdorferi* strains expressing OspA serotypes

The IPTG-inducible *mscarlet-I* viability reporter plasmid, pGW189 (34) was modified to facilitate the expression of *ospA* variants of interest under the regulatory control of the *P_ospAB_* promoter from *B. burgdorferi* strain B31. To accomplish this, transcription terminators were first introduced into pGW189 to prevent read through transcription from the *lacI* and *mScarlet*-*I* ORFs. The lambda_T0_rrnBT1 and rrnbT2 transcription terminators were amplified from plasmids pTN7 (49) and puc18_mtn7-lacZ_gentR(50) using NEB’s Q5 DNA polymerase and tailed primers listed in **Table S5.**

Amplified DNA fragments were then assembled into pGW189 immediately downstream of the *lacI* and *mscarlet*-I ORFs using the SacI/HindIII restriction sites and NEB’s HiFi Assembly kit, creating pGW206.

The *ospA* promoter and ORF from *B. burgdorferi* B31 were next amplified in two overlapping DNA fragments with tailed primers designed to introduce a silent SphI restriction site ∼50 bases into the *ospA* ORF and permit assembly into pGW206 via the PvuI and PvuII restriction sites.

The resulting viability reporter/OspA ST1 expression plasmid, pGW217, subsequently served as the basis for creating additional OspA expression plasmids. To accomplish this, pGW217 was first digested with SphI and PvuII to remove the *ospA*_ST1_ ORF immediately following the conserved lipoprotein insertion sequence (amino acid residue 17). Gene fragments from additional *ospA* serotypes/*Bbsl* genospecies were then synthesized (IDT) and assembled directly into the vector using HiFi DNA Assembly kit (NEB).

Silent mutations were introduced into *ospA* gene sequences when necessary to reduce sequence complexity for *in vitro* synthesis. Mutations were introduced at sites that resulted in a shift to the most common or second most common codon utilized by *B. burgdorferi* B31. The codon adaptive index for each *ospA* cDNA was assessed prior to synthesis using JCAT (51) to ensure that the introduced silent mutations did not reduce translation efficiency in *B. burgdorferi*.

Gene fragments were also synthesized to create *B. burgdorferi* B31, *B. valaisiana,* and *B. turdi* reporter plasmids with insertions, deletions, or substitutions at position 107 within the *ospA* ORF. To permit direct comparison with *B31* point mutants, a second “wild type” OspA ST1 reporter plasmid was also created containing the same silent mutations that were introduced to increase GC content/reduce sequence complexity for *in vitro* synthesis. The sequences of primers and synthesized *ospA* gene fragments used in this study are provided in **Tables S5, S6.** OspA reporter plasmids were sequenced and transformed into *B. burgdorferi* HB19-R1 (36), a high-passage derivative of the human blood isolate HB19 that lacks linear plasmid 54 (*ospAB*⁻), using previously described methods (52). Transformants were selected in liquid BSKII supplemented with gentamicin (100 µg/ml) and individual clones were isolated through serial dilution (52)

To evaluate native plasmid content in transformants, the plasmid profiles of *B. burgdorferi* HB19 and HB19-R1 were characterized using NEB’s Taq Quickload 2× PCR MM with established *B. burgdorferi* plasmid profiling primer sets (53). Additional primer sets targeting cp26, lp21, lp28-1, lp5, lp28-5, and lp28-6 were also included in the analysis (53, 54).Since plasmid content was uniform among clones transformed with mScarlet-I reporter plasmids carrying *ospA* ST1-7 (pGW217–pGW223, pGW250), plasmid profiling was not performed on HB19-R1 reporter strains encoding OspA ISTs due to the labor-intensive nature of the screening process. All *Bbsl* strains used or constructed in this study are listed in **Tables S1, S2.**

### Flow cytometry-based antibody binding to strains

To analyze the ability of bin 1 mAbs to bind to native OspA ST1-7 on the bacterial surface of primary isolates, *Borreliella* strains *B. burgdorferi* B31 (ST1), *B. afzelii* Pko (ST2), *B. garinii* PBr (ST3), *B. bavariensis* PBi (ST4), *B. garinii* PHei (ST5), *B. garinii* TN (ST6), and *B. garinii* T25 (ST7) (kindly provide by Dr. John Leong, Tufts University) were cultured at 33°C with 5% CO_2_ until mid-log phase. Bacterial cells were stored at −80°C in fresh media containing 20% glycerol, and preparation, incubations, and flow cytometry analysis was performed as described (46). To analyze the ability of bin 1 mAbs to bind to OspA ST1-7 on the bacterial surface of reporter strains HB19-R1, (GGW1072-1079; **Table S2**), bacterial cells were cultured at 33°C with 5% CO_2_ in BSKII media, minus gelatin and containing gentamycin 50 µg/ml, until mid-log phase. Bacterial cell preparation, incubations, and flow cytometry analysis was performed as described (55). The *Salmonella*-specific mAb, Sal4 IgG, was used as an isotype control (56).

### Complement-dependent borrelicidal assays

Complement-dependent bactericidal assays were performed as previously described (34) with several modifications. Frozen aliquots of OspA-expressing HB19-R1 reporter strains were thawed at room temperature and 500 µl from each stock (equivalent to 5×10^7^ spirochetes) were transferred to 50 ml conical tubes containing 45 ml of in gelatin-free BSK II supplemented with 50 µg/ml of gentamicin. Tubes were then sealed, and the cultures were incubated at 33°C under static growth conditions. Three days later, spirochetes were collected via centrifugation (4,000 x *g*), the supernatant was removed, andcell pellets were resuspended in phenol red-free BSK II with gentamicin (50 µg/ml) at a density of 3 x 10^7^ spirochetes/ml.

Spirochetes were next seeded into white bottom 96 well microtiter assay plates (Costar) and mixed 1:1 with serial dilutions of anti-OspA_ST1_ mAbs prepared in phenol red-free BSKII containing 5% guinea pig complement (Sigma-Aldrich) and gentamicin (50 µg/ml), resulting in 3×10^6^ spirochetes per well with a final complement concentration of 2.5% and antibody concentrations between 10.00 nM - 0.08 nM, or 20 nM - 0.16 nM (depending on strain/experiment). Eight untreated (antibody-free) control reactions per strain were also included on each assay plate to determine baseline and peak fluorescence for data normalization.

Following antibody addition, assay plates were incubated overnight at 37°C with 5% CO_2_ without agitation. The next day (18-20 h later), 1 mM IPTG was added to appropriate wells to induce expression of the mScarlet reporter in surviving spirochetes. Assay plates were then returned to the incubator for another 24 h period. The following day, median fluorescence intensity (MFI) was measured three times per plate at 569 nm (excitation) and 611 nm (emission) using a Spectromax ID3 plate reader (Molecular Biosystems) and Softmax pro version 7.1 software.

Mean MFI data was independently normalized for each assay plate using the untreated/non-induced and untreated/IPTG-induced controls to establish baseline (0) and peak (1) MFI. Graphpad Prism version 9.5.1 was then used to generate killing curves and heatmaps, while radar plots were constructed in Microsoft Excel. Ec_50_ values were assigned to the lowest antibody dilution resulting in >50% reduction in MFI relative to untreated controls. Ec_50_ values reported in heatmaps and radar plots reflect the mean calculated using normalized MFI data from three to five independent experiments. Strains that exhibited resistance to anti-OspA Bin 1 mAbs at the highest concentration tested (10 or 20 nM) were subjected to additional bactericidal assays with 10 µg/ml of the antibodies in question. Resistance at this dosage is shown as “>60 nm” in heat maps and radar plots.

### Cloning, expression and purification of OspA and anti-OspA Fabs

The PCR amplicon encoding *B. burgdorferi* OspA residues 18 to 273 was subcloned in frame into the pSUMO expression vector with an N-terminal deca-histidine and SUMO tag. All cloning was performed using a standard ligase independent cloning protocol. OspA was expressed in *E. coli* strain BL21 (DE3). The transformed bacteria were grown at 37°C in TB medium and induced at 20°C with 0.1 mM (IPTG) at an OD_600_ of 0.6 for ∼16 hours at 20°C. After induction, cells were harvested and resuspended in 20 mM Hepes pH 7.5 and 150 mM NaCl. The cell suspension was sonicated and centrifuged at 30,000 g for 30 minutes. After centrifugation, the protein-containing supernatant was purified by nickel-affinity and size-exclusion chromatography on an AKTAxpress system (GE Healthcare), which consisted of a 1mL nickel affinity column followed by a Superdex 200 16/60 gel filtration column. The elution buffer consisted of 0.5M imidazole in binding buffer, and the gel filtration buffer consisted of 20mM Hepes pH 7.5, 150mM NaCl, and 20mM imidazole. Fractions containing OspA (18-273) was pooled and subject to TEV protease cleavage (1:10 weight ratio) for 3 hours at room temperature in order to remove their respective fusion protein tags. The cleaved protein was passed over a 1mL Ni-NTA agarose (Qiagen) gravity column to remove the added TEV protease, cleaved residues, and uncleaved fusion protein. Each IgG was subjected to papain digestion followed by affinity depletion of the Fc fragment by Protein A FPLC chromatography to generate each Fab. After purification, each Fab and OspA were mixed in a 1:1 stoichiometry to form a stable complex concentrated to a final concentration of 10 mg/ml for all crystallization trials.

### Crystallization and data collection

Fab-OspA crystals were grown by sitting drop vapor diffusion at 4°C using a protein to reservoir volume ratio of 1:1 with total drop volumes of 0.2 μl. Crystals of each OspA-Fab complex were produced using the crystallization solutions shown in **Table S3**. All crystals were flash frozen in liquid nitrogen after a short soak in the appropriate crystallization buffers supplemented with 25% ethylene glycol. Data were collected at the 24-ID-E beamline at the Advanced Photon Source, Argonne National Labs. Data resolution limit was determined based on the I/s criterion which was 2.0 in the highest resolution shell. All data was indexed, merged, and scaled using HKL2000 (57) then converted to structure factor amplitudes using CCP4 (58).

### Structure determination and refinement

The structure of each Fab-OspA complex was solved by molecular replacement using Phaser (57). Molecular replacement calculations were performed using the Fab coordinates from PDB code 2XTJ as the search model for Fab 221-5, the Fab heavy chain coordinates from PDB code 4RIR along with the Fab light chain coordinates from PDB code 4M6O were independently used as search models for Fabs 221-11 and 857-2.

The Fab coordinates from PDB code 4M6O were also used as the search model for Fab 227-1. OspA coordinates (PDB code 1FJ1) were used as the search model for OspA in all four Fab-OspA complex structures solved. The resulting phase information from molecular replacement was used for manual model building of the Fab-OspA model using the graphics program COOT (59) and structural refinement employing the PHENIX package (60). Data collection and refinement statistics are listed in **Table S4**. Molecular graphics were prepared using PyMOL (Schrodinger) (DeLano Scientific LLC, Palo Alto, CA). The structures generated in this study were deposited in the Protein Data Bank (PDB; http://www.rcsb.org/pdb/) under accession numbers shown in **Table S4.**

### Phylogenetic tree construction

A collection of 135 OspA protein sequences with representatives from 23 recognized Bbsl genospecies was compiled from borreliabase.org, NCBI’s OspA IST typing database (PubMLST), and Bacterial and Viral Bioinformatics Resource Center (BV-BRC.org). MUSCLE (accessed via EMBL) was then used to generate a multiple sequence alignment of the collection. The resulting MSA file was then used to construct an unrooted approximately maximum likelihood phylogenetic tree using Fasttree and LG modelling (accessed via BV-BRC.ORG). The resulting phylogenetic tree was then formatted and visualized using ITOL version 7 (61). Files associated with phylogenetic tree construction are provided in **Dataset1**.

### OspA multiple sequence alignment (MSA)

A MSA was created containing one representative sequence from each of the 33 OspA clades/types identified through phylogenetic analysis. OspA variants selected for the alignment included OspA sequences utilized in viability reporter strain construction and OspA sequences from type or reference strains (when available). Sequence alignment was performed using the MAFFT algorithm with OspA from *B. burgdorferi* B31 as a reference. Following alignment, the Bin 1 epitope region (β strands 6-10, residues 82-146) was visualized using the MSA viewer interface on BV-BRC.org. A graphic of the alignment was then created with the Taylor color scheme and the sequence logo displayed to emphasize conservation between OspA types/clades.

### Microsphere immunoassay (MIA)

ST 1-7 and OspB were coupled to Magplex-C microspheres (5mg antigen/ 1×10^6^ microspheres) using a xMap Antibody Coupling Kit as recommended by the manufacturer (Luminex Corporation, Austin, TX). Successful coupling of ST 1-7 and OspB was confirmed using LA-2 and H6831, respectively. Beads were protected from light and stored 2-8°C in xMAP AbC Wash Buffer (5×10^6^ microspheres/mL) until use.

OspA bin 1 mAbs were serially diluted (1:2) in assay buffer (1 x PBS, 2% BSA, pH 7.4) in round bottom, non-treated plates (Costar, Kennebunk, Maine) with starting concentrations of 2.5 mg/mL (223–5), 0.625 mg/mL (857-2, 221-5, 221-20, 221-11, 227-1), or 0.3125 mg/mL (227-2).

Coupled microsphere stocks were diluted (1:50) in assay buffer and added (50 μL/well) to black, clear-bottomed, non-binding, chimney 96-well plates (Greiner Bio-One, Monroe, North Carolina). The mAb dilutions (50 μL) were combined with the microspheres and incubated at room temperature for 1 hr in a tabletop shaker (600 rpm). Plates were placed on a magnetic separator and washed three times using wash buffer (1 x PBS, 2% BSA, 0.02% TWEEN-20, 0.05% Sodium azide, pH 7.4). PE labeled goat anti-Human IgG Fc, eBioscience (Invitrogen, Carlsbad, California) secondary antibody diluted 1:500 in assay buffer was added (100 μL/well) and incubated at room temperature for 30 min in a tabletop shaker (600 rpm). Plates were washed as previously stated. The microspheres were resuspended in 100 μL of wash buffer and placed back on the tabletop shaker (600 rpm) for 5 minutes prior to analysis using a FlexMap 3D (Luminex Corporation).

## ACKNOWLEDGMENTS

We gratefully acknowledge Dr. John Leong (Tufts University) for providing *Borreliella* isolates and Dr. Meredith Finn (Moderna) for providing recombinant OspA proteins. We thank the Wadsworth Center’s Cell culture and media core for preparation of BSK II media, the Genomics core for DNA sequencing, and Drs. Renjie Song and Jennifer Yates of the Immunology Core facility for assistance in optimizing flow cytometry parameters.

## AUTHOR CONTRIBUTIONS

Conceptualization: GGW, MJR, NJM Methodology: GGW, MJR, CLP, GFG, LAC, DJV

Investigation: GGW, MJR, CLP, YC, GFG, DJV Visualization: GGW, MJR, CLP, GFG, DJV Supervision: MJR, LAC, NJM Writing—original draft: GGW, MJR, DJV, NJM

Writing—review & editing: GGW, MJR, DJV, NJM

## Data and materials availability

All data are available in the main text or the supplementary materials.

